# Rhometa: Population recombination rate estimation from metagenomic read datasets

**DOI:** 10.1101/2022.08.04.502887

**Authors:** Sidaswar Krishnan, Matthew Z. DeMaere, Dominik Beck, Martin Ostrowski, Justin R. Seymour, Aaron E. Darling

## Abstract

Bacterial evolution is influenced by the exchange of genetic information between species through a process referred to as recombination. The rate of recombination is a useful measure for the adaptive capacity of a bacterial population. We introduce Rhometa (https://github.com/sid-krish/Rhometa), a new software package to determine recombination rates from shotgun sequencing reads of metagenomes.It extends the composite likelihood approach for population recombination rate estimation and enables the analysis of modern short-read datasets. We evaluated Rhometa over a broad range of sequencing depths and complexities, using simulated and real experimental short-read data aligned to external reference genomes. In simulated datasets, the deviation from the expected value decreased as the number of genomes increased and we show that 80 genomes are sufficient to reduce these variations below 30%. Testing on an *S. pneumoniae* transformation experiment dataset we show that Rhometa accurately estimate the expected levels of recombination in a real world dataset.

## Introduction

A primary question in the field of microbial ecology is to understand the rate at which bacteria evolve and form species in nature (Didelot and Maiden 2010). A major driving factor of microbial evolution and speciation is recombination (Levin and Cornejo 2009; Didelot and Maiden 2010; Schmutzer and Barraclough 2019). Microbes are able to exchange and incorporate nucleotide sequences (DNA), genes or gene fragments, through homologous recombination, which often plays a greater role than *de novo* mutation for evolution (Rocha et al. 2005; Vos and Didelot 2009; Didelot and Maiden 2010; Iranzo et al. 2019). Furthermore, it is thought that recombination plays an important role in counteracting the effects of Muller’s ratchet, the theorised process where deleterious mutations inevitably accumulate over time leading to the loss of genomic fitness (Andersson and Hughes 1996; Vos 2009). Therefore, understanding the rate at which recombination occurs within a bacterial population can provide us insight into a crucial biological process that is necessary for their adaptation and survival.

The best way to study recombination in a microbial population is via metagenomics which allows us to study microbes in their natural environment via direct sequencing and analysis of environmental DNA (Thomas et al. 2012; Escobar-Zepeda et al. 2015). Shotgun metagenomic sequencing yields fragments of DNA sequences, referred to as reads, which taken together represent a random sampling of genome fragments from all the microbes in the environmental sample (Sharpton 2014). These reads can then be used to estimate the rates of recombination.

Within bacteria, recombination often takes the form of gene-conversion where homologous sequences of DNA are unidirectionally transferred from one cell and incorporated into another. This process can also occur between repeated sequences within the same bacterial chromosome and between homologous bacterial chromosomal sequences (Lassalle et al. 2015; Paulsson et al. 2017).

The rate of recombination within a population can be inferred using population genetic models for evolution. The Wright-Fisher model provides an analytical framework that quantifies various forces that can impact the evolution of a population such as random genetic drift and mutation (Tataru et al. 2017). Coalescent theory, building on the Wright-Fisher population model, provides an analytical framework for DNA polymorphism data and can be used to obtain quantitative estimates for recombination and mutation rates (Fu and Li 1999; McVean et al. 2002).

Coalescent theory provides the microbial (haploid) population scaled recombination rate, which is described as ρ = 2*N*_*e*_*r*, 2 × “effective population size” × “per individual” “per generation” “rate of initiation of gene conversion”, respectively (McVean et al. 2002) as well as the haploid population scaled mutation rate equation θ = 2*N*_*e*_*u*, 2 × “effective population size” × “per individual” “per generation” “mutation rate”, respectively (McVean et al. 2002). It is difficult to estimate *r* or *u* directly without additional prior information, so recombination and mutation rates are typically computed as the population scaled statistics ρ and θ or simultaneously as the ratio *r*/*u* also denoted as r/m (per site recombination to mutation rate) (McVean et al. 2002; Melendrez et al. 2016).

Several approaches have been used to estimate the recombination rate ρ. These include moment estimators, full-likelihood estimators and composite likelihood estimators. Moment estimators use summary statistics to estimate ρ, but their accuracy is limited by the fact that they cannot use all the genetic information available (Fearnhead and Donnelly 2001; Fearnhead and Donnelly 2002; Stumpf and McVean 2003). Full likelihood estimators are able to utilise all the genetic information available to them, but are so computationally intensive that their usage is impractical. To mitigate these issues and to make the approach more computationally tractable, composite likelihood estimators were developed (Hudson 2001; McVean et al. 2002; Stumpf and McVean 2003). With composite likelihood estimators, the scope of data that is analysed is reduced e.g. to only consider pairs of alleles, this approach is less computationally intensive with only a slight loss in accuracy compared to the full-likelihood approach (Hudson 2001; McVean et al. 2002; Stumpf and McVean 2003; Hermann et al. 2019).

There are several programs available that implement the composite likelihood approach for estimating the recombination rate, including LDhat (McVean et al. 2002; Auton and McVean 2007), LDhelmet (Chan et al. 2012), LDhot (Auton et al. 2014), PIIM (Johnson and Slatkin 2009) and Pyrho (Spence and Song 2019). Each are excellent for their respective use cases, but have limitations that make them unsuitable for modern read-based metagenomic datasets.

More specifically, LDhat (McVean et al. 2002; Auton and McVean 2007), LDhelmet (Chan et al. 2012) and LDhot (Auton et al. 2014) were designed for genome sequence analysis, not metagenomes. PIIM (Johnson and Slatkin 2009) was a pioneering attempt at a metagenomic read-based recombination rate estimator. While innovative at the time its application is impractical today. PIIM’s approach included computationally expensive techniques to integrate out uncertainty in low quality base-calls so as to retain as much information as possible from the scarce data available at the time. Today, deep sequencing is affordable and highly accurate, such that it’s often more practical to simply discard low quality sequence data rather than account for it computationally using complex algorithms. As such PIIM’s approach is impractical for the ever-larger datasets that are generated via modern sequencing techniques. Furthermore, it lacks support for modern sequence data formats (e.g. BAM), being limited to the obsolete ACE assembly format that is rarely used today.

Pyrho (Spence and Song 2019) is the latest composite likelihood estimator available at the time of writing and includes support for read based data in the form of VCF (variant call format) files, but this feature is only available for diploid organisms, which is not suitable for metagenomic (microbial/haploid) datasets where recombination occurs in the form of a gene-conversion process. Still other programs exist that calculate the population recombination rate through different approaches such as LDjump (Hermann et al. 2019) and CodABC (Arenas et al. 2015) which utilise summary statistics (Hermann et al. 2019), and programs such as ClonalFrameML (Didelot and Falush 2007; Didelot and Wilson 2015) which can provide an estimate of recombination rate relative to the mutation rate, but is designed around whole bacterial genomes.

Mcorr (Lin and Kussell 2019) is a program that can work with metagenomic reads and estimate the relative rate of recombination to mutation as well as the recombinational divergence, 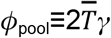, where 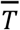 “is the mean pairwise coalescence time across all loci in the bulk pool” and *γ* the per base pair (bp) per generation recombination rate, equivalent to *r* in ρ = 2*N*_*e*_*r*. However, mcorr’s mathematical formulation is distinct from the well-known coalescent with recombination model and consequently from the population recombination rate (ρ = 2*N*_*e*_*r*), which may challenge interpretation. It is our aim to build on methods established in previous composite likelihood estimators for population recombination rate estimation to create a tailored solution that is applicable to modern read-based metagenomic datasets.

## New Approaches

Here, we present Rhometa, a software implementation of the composite likelihood based population recombination rate (ρ = 2*N*_*e*_*r*) estimation method, which builds upon the approach introduced in the LDhat pairwise program (McVean et al. 2002) that can be applied directly to modern aligned shotgun metagenomic read datasets. Details of its implementation are presented in the methods, while an evaluation of its accuracy on simulated and real data, and a comparison to existing tools are presented in the results.

## Results

### Simulated datasets

The development of our program was performed in two major phases, for the first phase we endeavoured to create a full genome recombination rate estimation pipeline for bacterial sequences based on the LDhat methodology (Rhometa_full_genome), once we were certain that we were able to replicate LDhat’s results exactly we then carefully adapted the program to work with read based datasets (Rhometa).

To evaluate LDhat and Rhometa_full_genome, we utilised msprime (Kelleher et al. 2016) to simulate bacterial sequences with recombination. Our simulations included multiple genomes (5-100 genomes) of size 25KB, under population recombination rates [5, 12.5, 25, 37.5, 50], recombination tract length 500bp, with 10 replicates (seed values 1-10) and population mutation rate 0.01. Lookup tables for population mutation rate 0.01 and population recombination rates 0-100 (101 steps) were used.

The LDhat pipeline configured for gene-conversion is available at: https://github.com/sid-krish/Nextflow_LDhat. Rhometa_full_genome pipeline is available at: https://github.com/sid-krish/Rhometa_Full_Genome. The full genome simulation pipeline is available at https://github.com/sid-krish/Nextflow_LDhat_Sim. A point of note is that the theta per site estimator is implemented separately by us as per equation 1 (McVean et al. 2002) in both our LDhat pipeline and Rhometa_full_genome pipeline, in contrast to the recombination estimation sections in LDhat. Additionally all variant sites are used for theta per site estimation, not just bi-allelic ones.

When simulating the population recombination rate with msprime, the number of samples (genomes), sequence length, gene conversion rate, gene conversion tract length, seed value and mutation rate are provided, the population_size was 1 (default) and the ploidy was fixed to 1. Default options are used in all other cases. The population recombination rate was calculated as such: 2 * ploidy * population_size * gene_conversion_rate * gene_conversion_tract_length. The recombination events were simulated first then mutation events were simulated on top, here the per site mutation_rate is provided and the population mutation rate per site was then calculated as such: 2 * ploidy * population_size * mutation_rate.

Initially the number of genomes was fixed and we varied the size of the genomes, but this analysis revealed that varying the genome size does not have a significant impact on the final population recombination rate estimations (Supplementary fig. S1). We therefore fixed the genome size and varied the number of genomes and in doing so we found that as the number of genomes increased the accuracy and variance of the final estimations also improved (fig. 1).

**FIG. 1.**
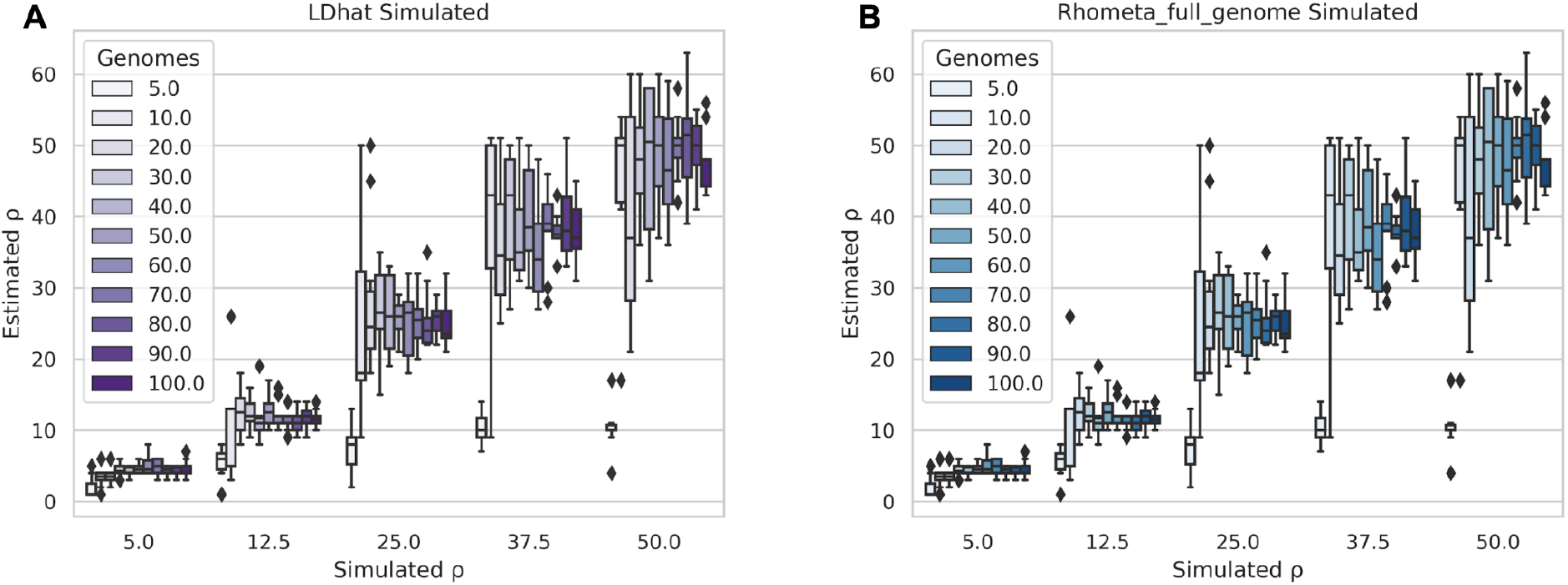
Comparison of LDhat and Rhometa_full_genome when running on simulated full genome. (**A**) LDhat. Simulated vs Estimated population recombination rate (ρ) for varying number of simulated full bacterial genome sequences. (**B**) Rhometa_full_genome. Simulated vs Estimated population recombination rate (ρ) for varying number of simulated full bacterial genome sequences.

We took a similar approach to evaluating the read based pipeline Rhometa to that used with LDhat and Rhometa_full_genome. For the read-based pipeline, the simulated full bacterial sequences, simulated via msprime, are further processed to be in the form of reads using the read simulator ART (Huang et al. 2012), these reads are then aligned to one of the bacterial sequences which represents the reference FASTA file, the first of the simulated sequences is used for this (fig. 4A). The aligned BAM and reference FASTA are then used for recombination rate estimation.

The simulation parameters were as follows: the population_size was 1 (default) and the ploidy was set to 1, number of genomes 20-200, genome size 100KB, population recombination rates [10.0, 20.0, 30.0, 40.0, 50.0], mean recombination tract length 1000bp, with 20 replicates (seed values 1-20) and population mutation rate of 0.01. Each seed value used applies to all aspects of the pipeline where a seed is required. The reads were paired-end of length 150bp, insert length 300bp, standard deviation of 25bp, with window size set to 1000 during analysis and the fold coverage values were [1, 4, 8, 16], with fold coverage ART program help defines it as such “the fold of read coverage to be simulated or number of reads/read pairs generated for each amplicon”. The lookup tables were generated and used for 3-250 (genomes), generated under population mutation rate 0.01 for population recombination rates 0-100 (0-1 in 101 steps plus 1-100 in 100 steps). Bam subsampling was also automatically applied by Rhometa during analysis if needed.

Additionally for the read-based pipeline, we evaluated the deviation of the estimated results from the simulated values. The formula used to calculate the deviation is (Estimated ρ (mean) - Simulated ρ) / Simulated ρ. This makes it easier to gauge the magnitude of deviation from the expected.

### Real Datasets - Transformation experiment

To further evaluate Rhometa we applied our pipeline on the data derived from a previously published laboratory transformation experiment, where the extent and distribution of recombination events were quantified. In the experiment (Croucher et al. 2012), *in vitro* recombination through transformation was performed on a *S. pneumoniae* strain. Transformed isolates were then sequenced and recombination events were identified. This dataset was also used to evaluate the mcorr method by its authors and as such it provides us with the opportunity to compare the results of our pipeline against those published in the mcorr paper.

The transformation experiments were performed with different concentrations of donor DNA, 5 ng mL^-1^ and 500 ng mL^-1^, 5 ng mL^-1^ and 500 ng mL^-1^ experiments had a similar number of recombination events, with the 5 ng mL^-1^ having a slightly larger number of events, the authors state that this indicates a single piece of DNA can act as the origin for multiple recombination events. The dataset is available in the form of reads, which Rhometa was designed to analyse. Each 5 ng mL^-1^ sample from experiment 1 was aligned to *S. pneumoniae* reference sequence ATCC 700669, NCBI accession NC_011900.1 the resulting BAM files were then merged and analysed with Rhometa.

To analyse the datasets, we first estimated the theta for median depth using the theta estimation pipeline, from which we obtained theta per site by default. We then generated lookup tables, based on the theta per site, for population recombination rates 0-20 in 201 steps for 3-200 genomes and used the lookup tables for the recombination rate estimation pipeline. Subsampling was enabled, with a window size of 1000 for paired end reads. As given in the Croucher *et al* paper, we used the value of 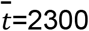 bp as the mean tract length for analysis. Additionally, we used 5 different seed values [0, 1, 2, 3, 4] for the subsampling step to account for any variance and then took the average of the values for recombination rate estimations.

Using the rho and theta estimates along with information from the experiment we also calculated the rho per site and r/m values. The default rho estimate, by Rhometa, is a whole-genome estimate. To obtain rho per site, the estimated rho value was divided by the tract length of 2300bp. To get the r/m value, we used the conversion formula (Didelot and Wilson, 2015): rho (per site)/theta (per site) * tract len * substitution probability. We calculated the substitution probability between the donor and recipient and found it to be (17534 − 385)/2221315 = 0. 00772, based on the information provided in the experiment paper (Croucher et al., 2012), where 17534 is the total reported number of variants called between donor reads & recipient genome, 385 is the number that is thought to be false positives, and 2221315 is the recipient genome size.

We repeated the process above for each 500 ng mL^-1^ sample from experiment 1 and the final merged BAM was analysed with Rhometa. Furthermore, the 5 ng mL^-1^ samples and 500 ng mL^-1^ samples in experiment 1 were analysed together, corresponding to 84 sequences. We prepared this dataset by merging the final 5ng and 500ng BAMs. We performed this analysis, to enable comparison with mcorr’s published results. The analysis was performed using the same process as with the 5 ng mL^-1^ samples and 500 ng mL^-1^ samples.

### Evaluation on Simulated Datasets

We first validated the full genome version of Rhometa (Rhometa_full_genome), which reimplements the core LDhat pairwise method to estimate rho. This was done to ensure accuracy in reimplantation of core LDhat algorithms which forms the basis for the read based (Rhometa) implementation. Comparison of estimated population recombination rate (rho) between LDhat and our reimplementation (Rhometa_full_genome), using our sweep of simulated genomes, shows identical results between LDhat (figure 1A) and our reimplementation (figure 1B), thus ensuring that we can captured LDhat’s algorithms accurately. With LDhat and our reimplementation, the number of genomes simulated has a large impact on the accuracy of the estimates, with results improving with higher numbers of genomes, especially at higher recombination rates.

We next evaluated Rhometa’s performance using our sweep of simulated read-sets. The number of simulated genomes had a large bearing on estimation accuracy (fig. 2), as also observed with LDhat, accuracy improved as the number of genomes increased and inter-replicate variance decreased as the coverage (fold_coverage) improved. This is especially evident for higher recombination rates. Larger population recombination rate values appear to require a relatively higher number of genomes for accurate estimation. For very low recombination rates between 0-1 (Supplementary fig. S3), the improvement in accuracy was not seen and a tendency to overestimate was observed.

**FIG. 2.**
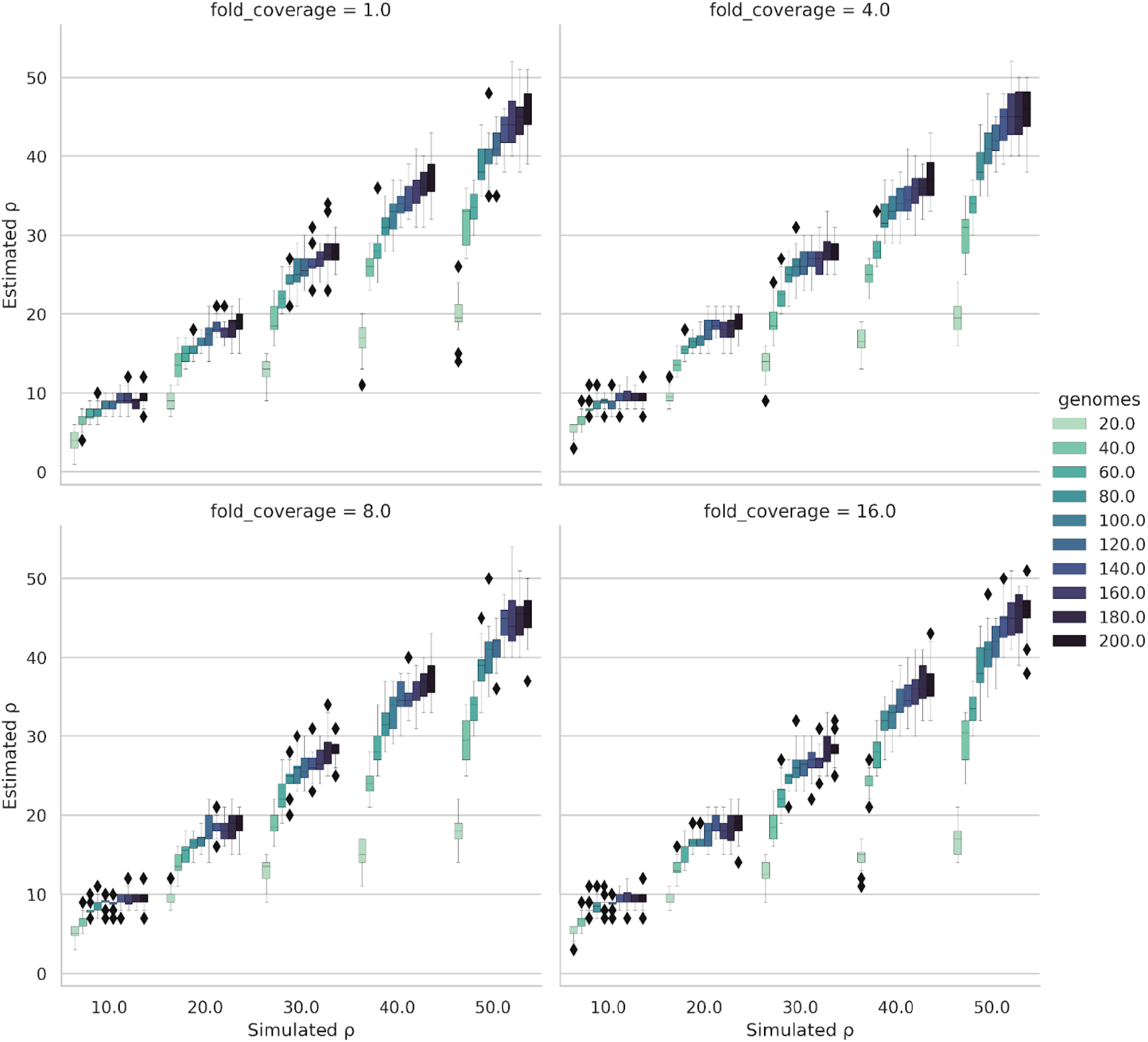
Simulated vs Estimated population recombination rate (ρ) results for Rhometa. Results for varying numbers of simulated genomes and fold coverage values for population recombination rates 10.0, 20.0, 30.0, 40.0, 50.0.

To get a clearer picture of the deviation of the estimated population recombination rate from the expected result, we calculated the deviation for the read based results (fig. 3). Here values closer to 0 indicated better performance, while values above 0 are overestimations, and values below 0 are underestimations (i.e. a deviation value of +/-0.1 would indicate that the final result is off by 10%). As the number of simulated genomes increased, the deviation of estimated to expected tended to decrease, achieving a deviation of less than 5-10% in most cases for a simulated rho of 50 with 200 genomes and 16x coverage. Such improvement is consistent with the patterns observed in LDhat. For simulated population recombination rates between 10-50, having greater than 80 genomes produced the least amount of deviation (generally within 20-30%), with the results significantly improving when more genomes are present. Our pipeline appears to be robust to variance in fold coverage. The differences between 16x coverage and 1x coverage being minor (fig. 3).

**FIG. 3.**
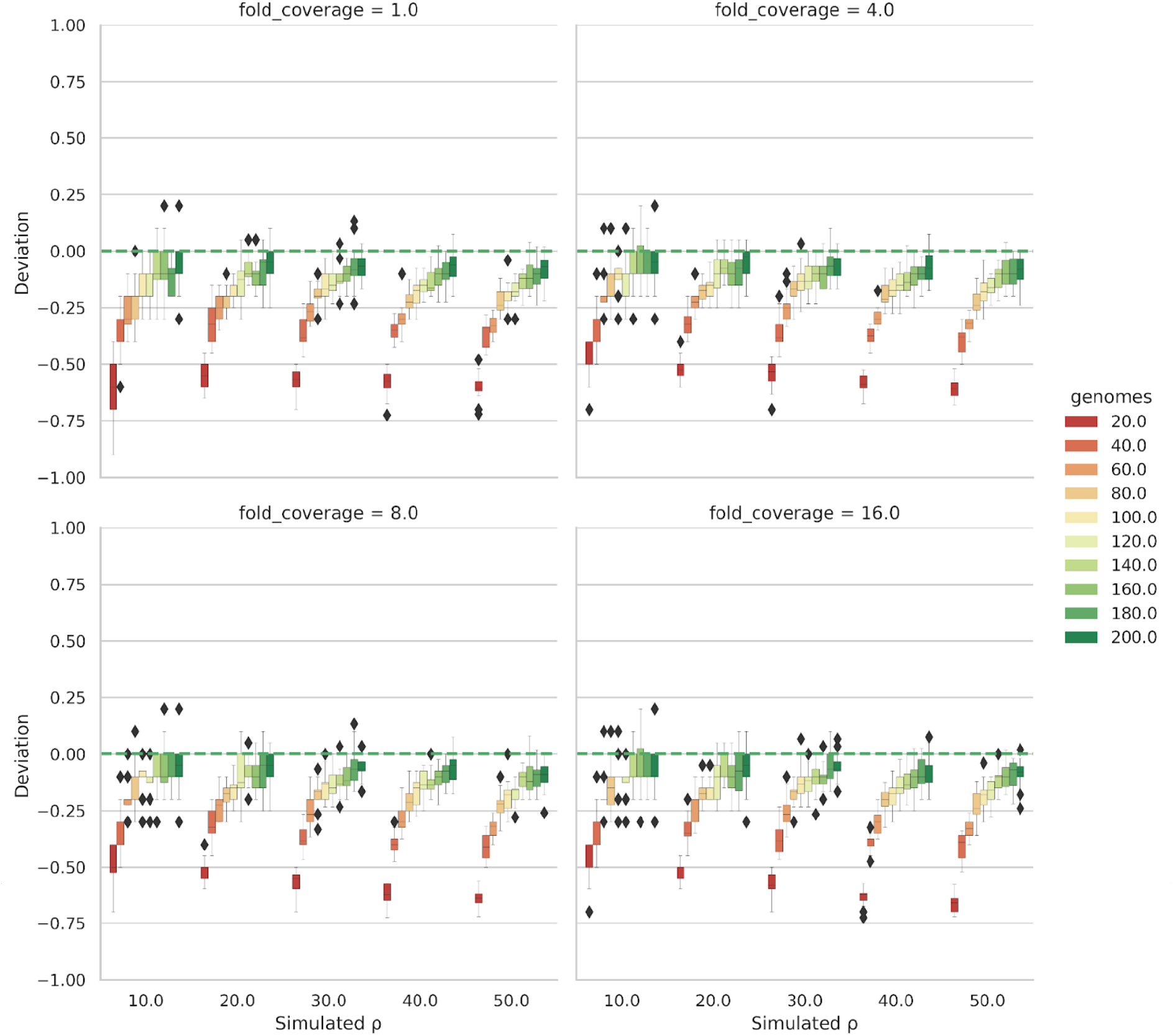
Deviation plot for results in fig. 2. Deviation is calculated as (Estimated ρ (median) - Simulated ρ) / Simulated ρ. Deviation results corresponding to figure 2 for population recombination rates 10.0, 20.0, 30.0, 40.0, 50.0.

### Evaluation on Real Datasets **-** Transformation Experiment

After establishing the performance of Rhometa on simulated datasets, we interrogated its performance on real short-read sequence data from a lab-based experiment designed to track and study recombination in *S. pneumoniae* (table 2). In the 5 ng experiment 1 dataset, we observed a seed averaged population recombination rate of 5.56, or a rho per site of 0.00242. Using median depth, the per site theta estimate was 1.8e-5, resulting in a per site ratio rho/theta of 134.4 and an r/m of 2386.4. Likewise for the 500 ng experiment 1 dataset, the population recombination rate was 5.22, or a rho per site of 0.00227, per site theta was 2e-5, the per site rho/theta per site was 113.5 and r/m was 2015.3.

**Table 1.** Pairwise table example.

|  | AA | AC | AG | AT | CA | CC | CG | CT | GA | GC | GG | GT | TA | TC | TG | TT |
| --- | --- | --- | --- | --- | --- | --- | --- | --- | --- | --- | --- | --- | --- | --- | --- | --- |
| (130, 136) | 0 | 13 | 0 | 21 | 0 | 0 | 0 | 10 | 0 | 0 | 0 | 0 | 0 | 0 | 0 | 0 |
| (130, 143) | 13 | 0 | 21 | 0 | 0 | 0 | 10 | 0 | 0 | 0 | 0 | 0 | 0 | 0 | 0 | 0 |
| (130, 169) | 0 | 29 | 1 | 3 | 0 | 8 | 0 | 0 | 0 | 0 | 0 | 0 | 0 | 0 | 0 | 0 |
| (130, 311) | 2 | 0 | 26 | 0 | 0 | 0 | 9 | 0 | 0 | 0 | 0 | 0 | 0 | 0 | 0 | 0 |
| (130, 358) | 0 | 0 | 0 | 19 | 0 | 2 | 0 | 6 | 0 | 0 | 0 | 0 | 0 | 0 | 0 | 0 |

Note - The coordinates of variant site pairs are relative to the reference genome, while bases at these sites are taken from the reads.

**Table 2.** *S. pneumoniae* transformation experiment analysis.

|  | Rho | Rho<br>(per site) | Theta<br>(per site) | Rho/theta<br>(per site) | r/m |
| --- | --- | --- | --- | --- | --- |
| 5 ng mL<sup>-1</sup> | 5.56 | 0.00242 | 1.8e-5 | 134.4 | 2386.4 |
| 500 ng mL<sup>-1</sup> | 5.22 | 0.00227 | 2e-5 | 113.5 | 2015.3 |
| 84 sequences | 4.48 | 0.00195 | 1.8e-5 | 108.3 | 1922.97 |

Note - The results from using Rhometa to estimate mutation and recombination rates on an *S. pneumoniae* transformation experiment. Rho and Theta (per site) are estimations from the program, whereas the others values are derived. The rows represent different datasets corresponding to concentrations of donor DNA, 5 ng mL<sup>-1</sup> and 500 ng mL<sup>-1</sup>, which were studied as part of the experiment. 84 sequences represent the combined 5 ng mL<sup>-1</sup> and 500 ng mL<sup>-1</sup> datasets. Rho is the population recombination rate for the whole genome, Rho (per site) is derived by dividing by the tract length (2300), Theta (per site) is a per site population mutation rate estimate and the Rho/theta (per site) is a ratio of the two. r/m is per site recombination to mutation rate.

For the dataset which combines all 5 ng and 500 ng experiment in one file (84 sequences), the population recombination rate was 4.48, or a rho per site of 0.00195, per site theta was 1.8e-5, the per site rho/theta was 108.3 and the r/m was 1922.97.

To compare the results for the same transformation experiment with those of mcorr, the estimated r/m values were used. The authors of mcorr provide **γ**/**μ** (similar to r/m) for the evolved strain (reads) representing combined 5 ng mL^-1^ samples and 500 ng mL^-1^ samples in experiment 1 (84 sequences) where they estimate a **γ**/**μ** value of 0.93. Due to the nature of the experiment it was expected that the rate of recombination would be far higher than the rate of mutation. As an experiment designed to induce transformation over short timescales, this should lead to a large excess of substitutions derived from recombination events, relative to *de novo* mutation. As such, mcorr potentially significantly underestimated the true value and Rhometa better reflects our *a priori* expectation for the 84 sequence dataset with an estimated r/m ratio of 1922.97.

Based on the information provided in the Croucher *et al* (Croucher et al. 2012), the average number of bases changed by recombination (240) and mutation (0.631), we can calculate the average r/m for a genome to be 240/0.631 = 380.3, for the 84 sequence dataset. The actual value should be higher due to the fact that a region can experience multiple recombination events, so this would be a lower bound estimate. With Rhometa we do observe a value greater than 380.3 of 1922.97 for the 84 sequence dataset. The values for the average number of bases changed by recombination and mutation are derived as follows. In the Croucher *et al* paper, they state that the mean proportion in the recipient genome changed due to recombination was 1.4% we estimated the average number of bases changed by recombination in a single genome as: 2221315 (recipient genome size) * 0.014 * 0.00772 (substitution probability) = 240.08. Additionally, in the paper it is stated that there were 2,312 polymorphic sites, 59 of which not coming from the donor, 6 of these sites were false positives, with the others likely being *de novo* point mutations or intragenomic recombinations. We take the upper bound for *de novo* mutations to be 53, then to get the average number of *de novo* mutations per genome we can divide by 84, the total number of sequences in the combined 5 and 500 ng dataset, 53/84=0.631.

## Discussion

Recombination plays a crucial role in microbial evolution and speciation (Levin and Cornejo 2009; Didelot and Maiden 2010; Schmutzer and Barraclough 2019). Understanding the rate at which recombination occurs provides us an insight into the impact of this process. Metagenomics is the only method that allows us to study recombination in real-world natural microbial communities without culture bias (Wooley et al. 2010). However, there are currently no software tools to accurately estimate population recombination rates on large metagenomic datasets. To fill this gap, we have developed Rhometa, a software implementation that builds on the well known composite likelihood estimator for population recombination rate estimation and enables interrogation of next generation sequencing reads from shotgun metagenomic experiments.

Rhometa enables the analysis of modern short-read metagenomic datasets to accurately quantify population recombination rates for naturally occurring prokaryotic populations. Composite likelihood population recombination rate estimators are among the most accurate methods known, and our implementation makes these methods available to the wider metagenomics community. This is significant as most microbes cannot readily be cultured, by some estimates only 1-15% are readily cultivable in laboratories (Singh et al. 2009).

Shotgun metagenomics yields reads from microbes taken directly from the natural environment and mitigates issues related to culture dependent studies. However, there has until now not been a viable approach for quantifying the population recombination rate from these reads. PIIM (Johnson and Slatkin 2009) and mcorr (Lin and Kussell 2019) come closest to being applicable to shotgun metagenomic datasets, being designed for this use case. However in the case of PIIM its statistical model uses a very compute intensive approach to account for low quality data, making the compute requirements impractical for large modern datasets. Meanwhile, with mcorr, the mathematical formulation is distinct from the well-known population recombination rate (ρ = 2*N*_*e*_*r*), which may represent challenges for interpretation. On an experimental dataset where transformation was used to produce a population of recombinants for sequencing, the approach implemented in mcorr appears to significantly underestimate the recombination rate.

Rhometa is well positioned to exploit the abundance of preexisting metagenomic datasets to enable a thorough first-pass study of recombination rates in microbial communities.

Rhometa represents a viable solution for population recombination rate estimation from next generation sequencing based datasets including gene-conversion based recombination. Using simulated and experimental datasets we demonstrated that our implementation accurately detected population recombination rates.

To build our program, our approach was to first reimplement the LDhat pairwise program for gene conversion. Doing so we were able verify that we accurately captured the core algorithms of LDhat while having a modern and adaptable implementation around it. The simulation results in figure 1 A,B for LDhat and the Rhometa_full_genome respectively show that we were able to reproduce the LDhat results 1:1. Having validated the reimplementation we then adapted it for read based datasets. LDhat is effective at detecting changes in the magnitude of the simulated population recombination rates, and produces accurate estimates for cases with large numbers of genomes (fig. 1A). Our analysis showed a trend where the accuracy and variance of the estimates improved as the number of genomes increased.

We then evaluated the performance of Rhometa on simulated datasets and the results (fig. 2 and 3) demonstrated that the read-based pipeline performs well and consequently represents a successful implementation of the composite likelihood population recombination rate estimator for metagenomic read-based datasets. As with LDhat and Rhometa_full_genome, the performance of the read based pipeline improves with the number of genomes present, having 80 genomes or more produces the best results. Very small rho values, those between 0-1 (supplementary fig. S3), are an exception as the implementation has a tendency towards over estimation.

Rhometa was further applied to a *S*.*pneumoniae* transformation experiment (Croucher et al. 2012), where the extent of recombination could be directly quantified. This dataset was also analysed by the authors of mcorr for their paper. The transformation experiment used different quantities of donor DNA, 5 ng mL^-1^ and 500 ng mL^-1^. When comparing the results of our pipeline with mcorr, using a combined 5ng and 500ng dataset representing 84 sequences, as was done with mcorr, Rhometa was able to accurately detect the higher rate of recombination relative to mutation as expected. From direct evidence we estimated a conservative lower bound for the ratio of recombination to mutation as r/m > 380.3 for the 84 sequence datasets. Rhometa was able to meet this condition by calculating an r/m value of 1922.97 for the 84 sequence dataset, while mcorr estimates a r/m value of 0.93. A point of note is that r/m for the 84 sequence datasets is lower than the individual 5 ng and 500 ng results (table 2), this lower r/m value may be an artefact of combining both 5 ng and the 500 ng datasets.

The main difference between the LDhat approach and the read based approach is as follows. In LDhat the final population recombination rate estimate “is obtained by combining the likelihoods from all pairwise comparisons” (McVean et al. 2002), the likelihoods come from the pregenerated lookup tables as mentioned in methods. For genome sequence datasets this means we can use a likelihood table generated for the exact number of genomes/depth in a dataset, the number of genomes represents the depth which is fixed and all pairs of sites we look at will have this depth. On the other hand, for aligned read based datasets the main complication is that the depth can vary greatly from site to site. We addressed this issue by using an appropriate depth lookup table for the variant site pairs being considered, the rationale for which is that likelihood for the variant site pairs considered is obtained individually and then combined for a final composite likelihood. Taking into account that the likelihoods are obtained individually for each pair of variant sites, we bin variant site pairs based on depth, and then use the appropriate depth likelihood table to obtain the likelihood for each pair and then finally combine the likelihoods to get a result for the entire dataset. Additionally, we have introduced a novel weighted sum when calculating the composite likelihood across coverage depth (equation 2). Rhometa thus enables the application of the composite likelihood estimator approach for current shotgun metagenomic datasets.

An important advantage of Rhometa and its use of raw reads over a consensus assembly from each sample, is that the potential microdiversity within each dataset is preserved for analysis.

Another advantage of our pipeline is that when preparing a metagenomic dataset for analysis with Rhometa, very little preprocessing is required. Short reads can be aligned to existing reference genomes for a species or to a reference MAG (Metagenome-Assembled Genome). The BAM file from the alignment and the reference genome/s used in the form of FASTA is all that is needed. As discussed, Rhometa performs better the more genomes there are, it is possible to get a minimum count for the number of genomes present when simulated under the coalescent model with recombination. In real metagenomic samples, any single sample may have millions of genomes of the same species, and across samples there may be significant population structure that is not captured by the standard coalescent model with recombination. The relationship between the number of metagenomic samples, the depth of sequencing of each sample, and the genome count in our simulation study is therefore not straightforward.

### Limitations and Future Directions

While we have endeavoured to make a complete package with Rhometa that addresses all aspects of population recombination rate, there are a few limitations. One such limitation is the automatic inference of tract length, which is also not possible with LDhat(McVean et al. 2002) or PIIM (Johnson and Slatkin 2009). In the context of the composite likelihood approach, the authors of both LDhat and PIIM suggest that while it may be theoretically possible to co-estimate the population recombination rate and tract length, in practice it is challenging. Instead, following the examples of LDhat and PIIM, Rhometa fixes the average tract length for population recombination rate estimation. As observed by the authors of PIIM, tract length tends to rescale the population recombination rate estimate and large mispecifications can cause further deviations (Johnson and Slatkin 2009).

Furthermore, the nature of our method is not sensitive to very low rates of recombination as observed when attempting to evaluate rates between 0-1 and we suggest exercising caution for such fine scale analysis.

Another point of note concerns the generation of the lookup tables for the program. While it is relatively fast to generate lookup tables due to the incorporation of LDpop, it can still require substantial time for a high-resolution table with a large number of samples.

Generation of lookup tables require specification of theta per site, however in our tests we have found that for realistic values of theta per site (i.e. less than 0.01), where 1% of sites have experienced mutation, estimation of the population recombination rate is relatively insensitive to changes in theta per site.

We believe the availability of a tool such as Rhometa, which can be easily applied to current metagenomic datasets is timely and significantly expands the range of habitats and therefore microbial communities that can be studied for recombination, giving us an insight into the extent to which they can adapt and speciate. How rho varies within environments and between taxa is unknown, Rhometa can help investigate many such fundamental questions related to the evolution and survival capacity of microbes. With the aid of data analysis techniques, metagenomic datasets can be further combined with environmental and sequencing metadata to help study the intricacies of recombination. Many ecological factors can modulate and effect recombination (González-Torres et al. 2019). Synthesis of other data types with the results of our program may yield a clearer understanding of such relationships. We have built our approach in a modular and easy to adapt manner making this and similar applications easy to explore in the future.

## Methods

Our approach focuses on advancing the composite likelihood approach for use with metagenomic read datasets. We have built our metagenomic population recombination rate estimator program upon the approach introduced in the LDhat program, specifically the LDhat pairwise module (McVean et al. 2002). LDhat is a well known and used program with support for microbial datasets, specifically for the gene-conversion type recombination which occurs in microbes, however, it is limited to aligned genome sequences. We have carefully adapted the program to work with modern read based metagenomic datasets. Additionally for our implementation, we have subsumed features from Pyrho (Spence and Song 2019). Pyrho, while lacking support for microbial (haploid) datasets, is a modern composite likelihood estimator implemented in python. Like Pyrho, our program is also implemented in python and aims to make use of modern libraries and their features. As a result of this shared implementation approach, we were able to call applicable functionalities from Pyrho, helping avoid unnecessary code rewrites.

### User Input

As input, the Rhometa pipeline requires a FASTA format reference sequence and a BAM file of metagenomic reads of interest aligned to the reference. In our pipeline, we have used BWA MEM (default parameters) to produce the input BAM file (Li 2013).

### Variant site pairs

The first step of the pipeline involves identifying variant sites (also known as segregating sites). Our program first filters the user supplied BAM for mapping quality and relative alignment score and subsequently performs variant calling against the user supplied reference FASTA using the program freebayes (default parameters with -p (ploidy) = 1) (Garrison and Marth 2012). The resulting VCF file, containing information on all predicted variant sites, is reduced to only single nucleotide polymorphisms (SNPs) using bcftools (Danecek et al. 2021).

Rather than individual variant sites, the composite likelihood estimator as implemented in LDhat considers variant site pairs, tracking count and position within the reference genome’s coordinate space to estimate the recombination rate. For instance, if variant sites are found at reference positions 1, 3, and 5 the set of variant site pairs would then be (1, 3), (1, 5), and (3, 5).

### Pairwise table

The LDhat pairwise module was designed for genome sequences and considers all possible variant site pair combinations across the sequences being analysed. Rhometa restricts its consideration to the set of variant site pairs linked by individual reads or read-pairs. For single-end reads, both sites within a variant pair must fall within the extent of an individual read, while for paired-end reads variants can fall within the insert length. A separation limit of 1000 bp is imposed on paired end variant site pairs reflecting a practical upper limit on insert size for current Illumina short-read sequencing technology (Tan et al. 2019).Rhometa performs well with both single and pair-end reads, with very little difference in the results between the two (supplementary fig. S2)

For all accepted variant site pairs, we construct a pairwise table of observational frequency (table 1). The pairwise table allows for the possibility of all 16 combinations for any variant site pair. The table also captures the fact that multiple reads can align at a position. Instances where variant site pairs contain an ambiguous base (eg. N) are ignored.

### Splitting the pairwise table by depth

In the pairwise table, variant site pair total alignment depth is calculated by row summation (e.g. For the pair (130, 136) from (table 1), total depth is 13 + 21 + 10 = 44. For the whole genome approach of LDhat, this marginal value is a constant, while for metagenomic data depth of coverage can vary greatly across sites. As such it is necessary to split the pairwise table into constant depth subtables so that the depth can be taken into account and handled in downstream processing.

### Bi-allelic pairwise table

For each constant-depth subtable, sites that do not contain two alleles are excluded (only biallelic sites should be in the pairwise table).

### Lookup tables

Lookup tables improve the computational efficiency of the composite likelihood approach by precomputing the likelihoods for different configurations of sets of allele pairs. Lookup tables are generated under a fixed population mutation rate and a range of population recombination rates, typically between 0 - 100 (McVean et al. 2002; Auton and McVean 2007). We use the program LDpop (Kamm et al. 2016) for generating lookup tables as it is the most feature rich and most efficient program of its kind currently. Details on how the lookup tables are used can be found in Appendix A. It is a standard process for which we have made use of some functions from Pyrho to avoid reimplementation. Generation of lookup tables with ldpop are parameterised by the number of genomes, range of population recombination rates, and theta per site. Further, the “approx” option is used which is significantly faster but still quite accurate when compared to the ldpop’s exact algorithm.

### Watterson’s theta estimate

A subprogram is provided to estimate the population mutation rate (θ = 2*N*_*e*_*u*), per site, based on Watterson’s theta estimate as implemented in LDhat. The program requires the aligned BAM file and the reference FASTA file and makes use of freebayes to identify variant sites which is required for the Watterson estimate. Theta per site is a required parameter for lookup table generation adjusted for read based datasets the theta per site estimate is calculated based on dataset depth – specifically mean and median depth – in place of the number of sequences (supplementary fig. S4).

### Lookup table and depth

The number of alignments covering a variant site pair (the depth) determines which constant-depth lookup table to use for precomputed likelihoods. In cases where high depth values are not covered by the generated lookup tables, a subsampling feature is included that is able to downsample the BAM to a given depth. This ensures that positions with a depth exceeding that of the highest generated lookup table are not omitted from consideration. BAM subsampling uses a random sampling process and permits a list of seed values for testing and identifying any variance that can stem from the subsampling. In general, if the depth of the largest available lookup table is small, an increased need for downsampling could result in a decrease in estimation accuracy.

### Calculating r_ij_

The next step is to calculate recombination rate values for each variant site pairs, these values are denoted by r_ij_, with i and j referring to the variant sites. The method used for calculation differs for crossing-over and gene-conversion modes of recombination. Microbes undergo recombination via gene-conversion and the equation used to calculate r_ij_ is as follows (McVean et al. 2002):

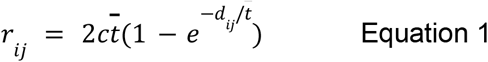

For equation 1, *c* represents the per base rate of initiation of gene conversion, 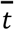 the average gene conversion tract length and *d*_*ij*_ the distance between a variant site pair. 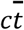 is taken together and represents the range of population recombination rates being evaluated, this is typically between 0 - 100 and is the same as the range of rho values used when generating the lookup tables. The process essentially involves computing *r*_*ij*_ for each variant site pair for the range of population recombination rate values.

### Final pairwise likelihoods

Next, we bring together the information we have generated thus far: the matched likelihoods for the variant site pairs and the *r*_*ij*_ values table from the previous step. For each variant site pair, we use the matched likelihood values for recombination rates, and on these apply linear interpolation to determine the likelihood value for *r*_*ij*_ value for that variant pair configuration, the *r*_*ij*_ is compared against the range of recombination rates in the matched likelihoods table. This process is done for all the variant site pairs and the resulting likelihoods table are the final likelihoods for a given range of population recombination rates being evaluated, which again is typically between 0 - 100.

### Population recombination rate

As the number of observations provided at a given depth represents the degree of evidential support towards the final rho estimate, we introduce a novel weighting algorithm biased towards higher depth and observation count as follows:

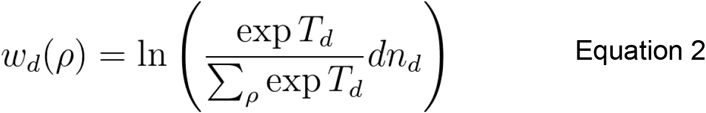

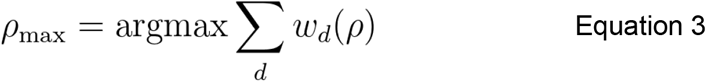

Here, *d* represents depths observed in the dataset. Weighting is performed on each per-depth table denoted by *w*_*d*_(ρ) (equation 2). In the right side of the equation *T*_*d*_ is the unweighted per-depth table and *n* the number of unique variant site pairs. The reweighted log-likelihoods *w*_*d*_(ρ) are summed and the maximum log likelihood value (closest to 0) corresponds to the most likely final estimated population recombination rate *ρ*_*max*_ (equation 3).

### Program Structure

The program is organised into 4 pipelines, each dedicated to a specific task. These pipelines are written using nextflow, a framework for pipeline management (Di Tommaso et al. 2017). All the scripts used in the individual pipeline steps were written using the python programming language and various python libraries. Some python scripts were adapted or used as is from the programs LDpop (Kamm et al. 2016) and Pyrho (Spence and Song 2019). Additional programs used in the pipelines include msprime (Kelleher et al. 2016), ART (Huang et al. 2012), BWA MEM (Li 2013) and samtools (Li et al. 2009).

The four pipelines, sim_gen, theta_est, lookup_table_gen and rho_est (fig. 4), correspond to the nextflow pipeline names, e.g. sim_gen.nf within Rhometa, and perform the functions defined in the following paragraphs.

**FIG. 4.**
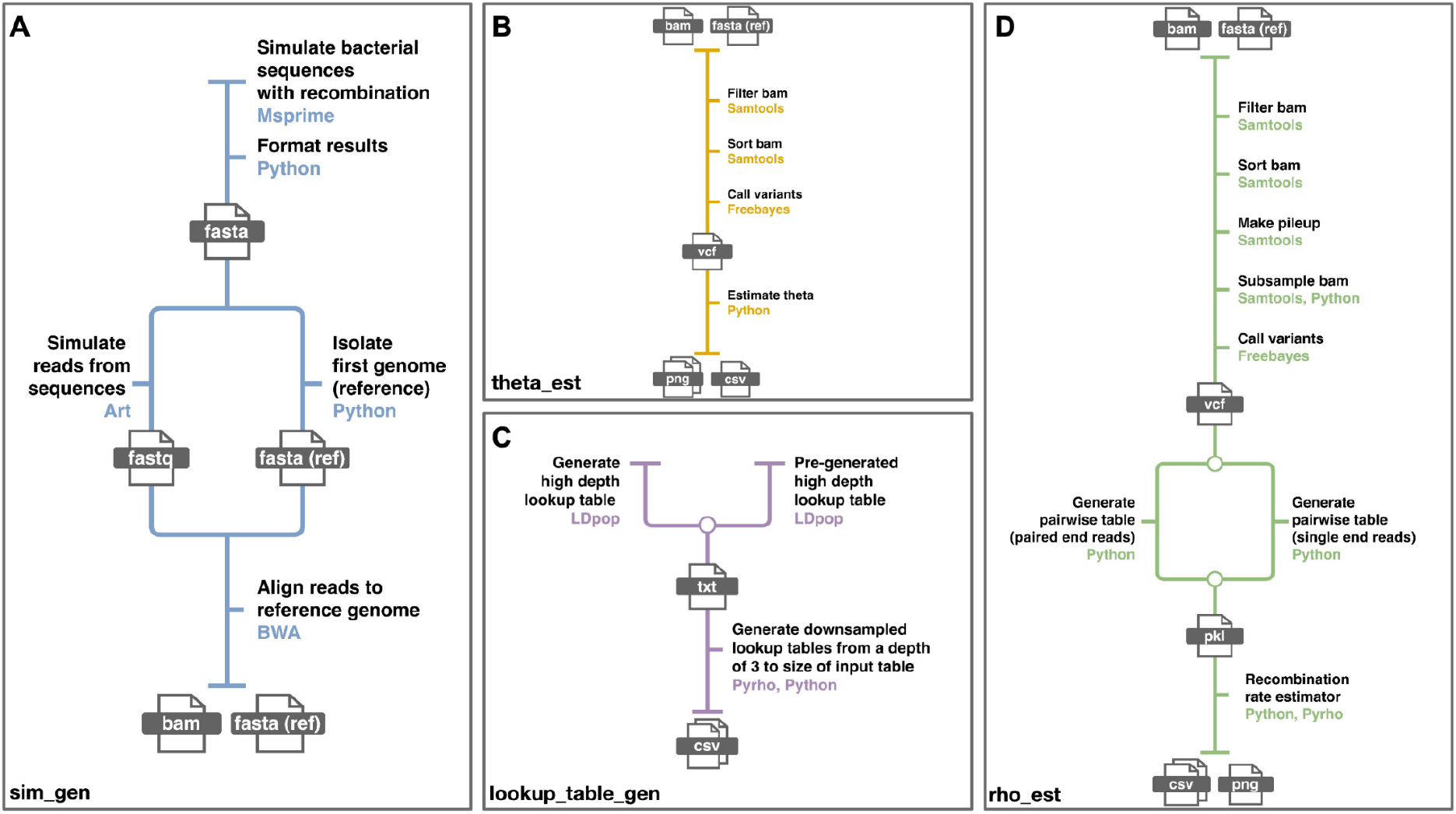
The pipelines that together make up Rhometa. (**A**) Pipeline for generating simulated metagenomic read datasets. (**B**) Pipeline for estimating the population mutation rate (**C**) Pipeline for generating the lookup tables required for the recombination rate estimator (**D**) Pipeline for estimating the population recombination rate.

Sim_gen (fig. 4A) is used to generate BAM files and FASTA reference files with simulated reads from bacterial genomes with recombination, the bacterial genomes were simulated using msprime. This pipeline is primarily included so that the simulated datasets used for this paper can be reproduced, but is not required to analyse real datasets.It is in a separate repository and can be accessed at: https://github.com/sid-krish/Rhometa_sim

Theta_est (fig. 4B) is used to determine the population mutation rate (theta) per site based on the Watterson estimate as implemented in LDhat, details in methods. This pipeline estimates theta on the dataset of interest, furthermore, theta per site is one of the required parameters for generating lookup tables. The user has the option to use the estimated theta or a different value when generating lookup tables.

The Lookup_table_gen (fig. 4C) component of the pipeline makes use of LDpop and Pyrho to generate the lookup tables required for the recombination rate estimator and can be launched in one of 2 ways. It can either use a pre-generated lookup table for high depth, which then will be downsampled for each depth from 3 to the depth of the lookup table or the pipeline can generate a high depth lookup table from scratch and then perform the downsampling step. The downsampling algorithm is a part of Pyrho, it is significantly faster to generate the required smaller lookup tables from a larger table via downsampling and the results are essentially identical.

The rho_est pipeline (fig. 4D) is used to estimate the population recombination rate of metagenomic read based datasets provided in the form of BAM and reference FASTA files. It makes use of the lookup tables generated by the lookup_table_gen pipeline.

Rhometa is available at: https://github.com/sid-krish/Rhometa

Our pipelines for evaluating LDHat, the Rhometa_full_genome pipeline and the simulated dataset generator for these pipelines can be accessed here:

- LDhat Nextflow Pipeline: https://github.com/sid-krish/Nextflow_LDhat
- Rhometa Full Genome Pipeline: https://github.com/sid-krish/Rhometa_Full_Genome
- Nextflow_LDhat_sim (used for both Rhometa Full Genome and LDhat Nextflow Pipeline) : https://github.com/sid-krish/Nextflow_LDhat_Sim

## Data Availability

https://doi.org/10.26195/0w2e-tt98

## Acknowledgements

This work was supported by an Australian Government Research Training Program Scholarship. This research was supported by the Australian Government through the Australian Research Council Discovery Projects funding scheme under the project DP180101506, http://purl.org/au-research/grants/arc/DP180101506 (to AED). The funders had no role in study design, data collection and analysis, decision to publish, or preparation of the manuscript.

## Supplementary

**Figure S1.**
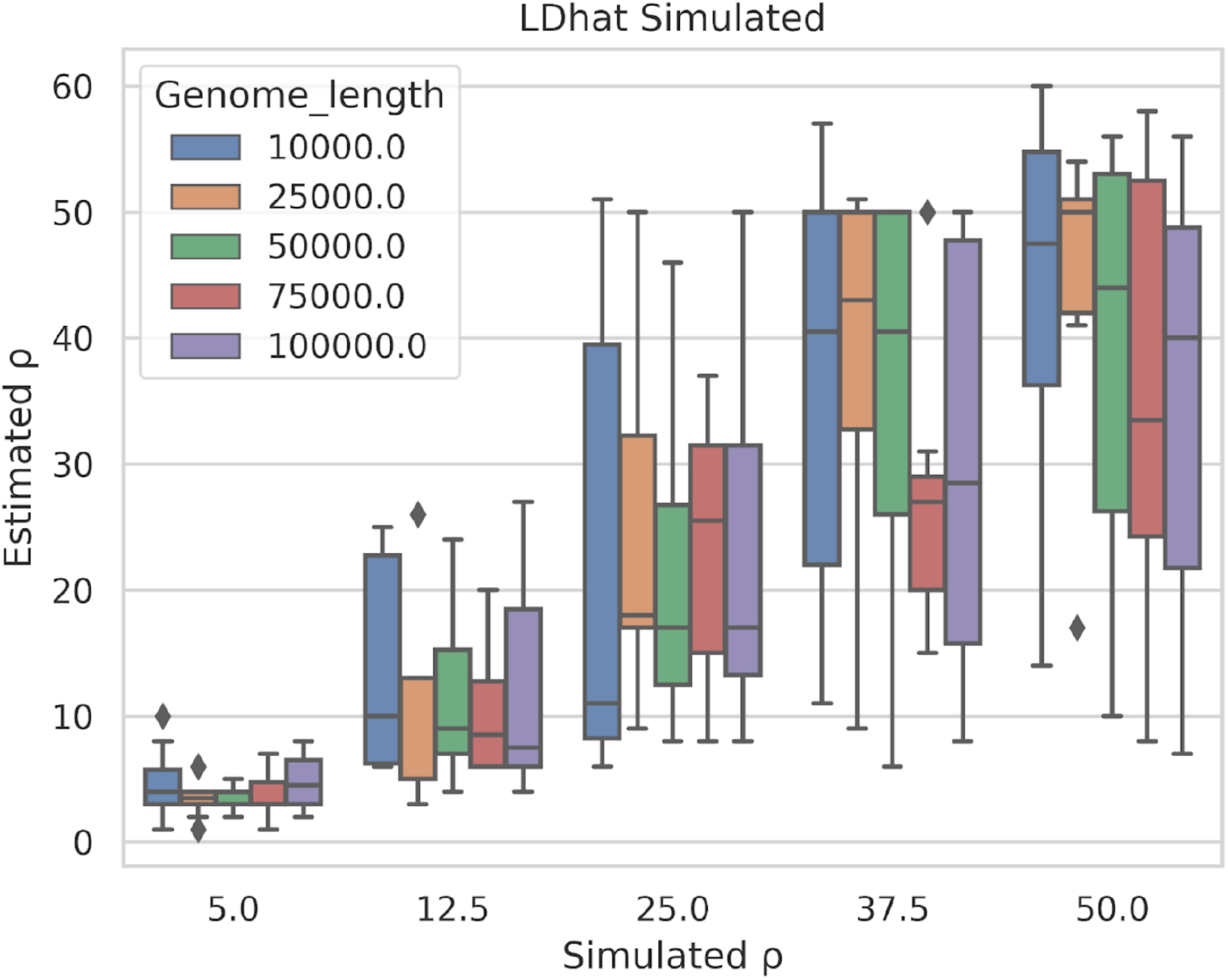
Results of varying simulated genome lengths for testing LDhat (number of genomes fixed at 10, tract length 500)

**Figure S2.**
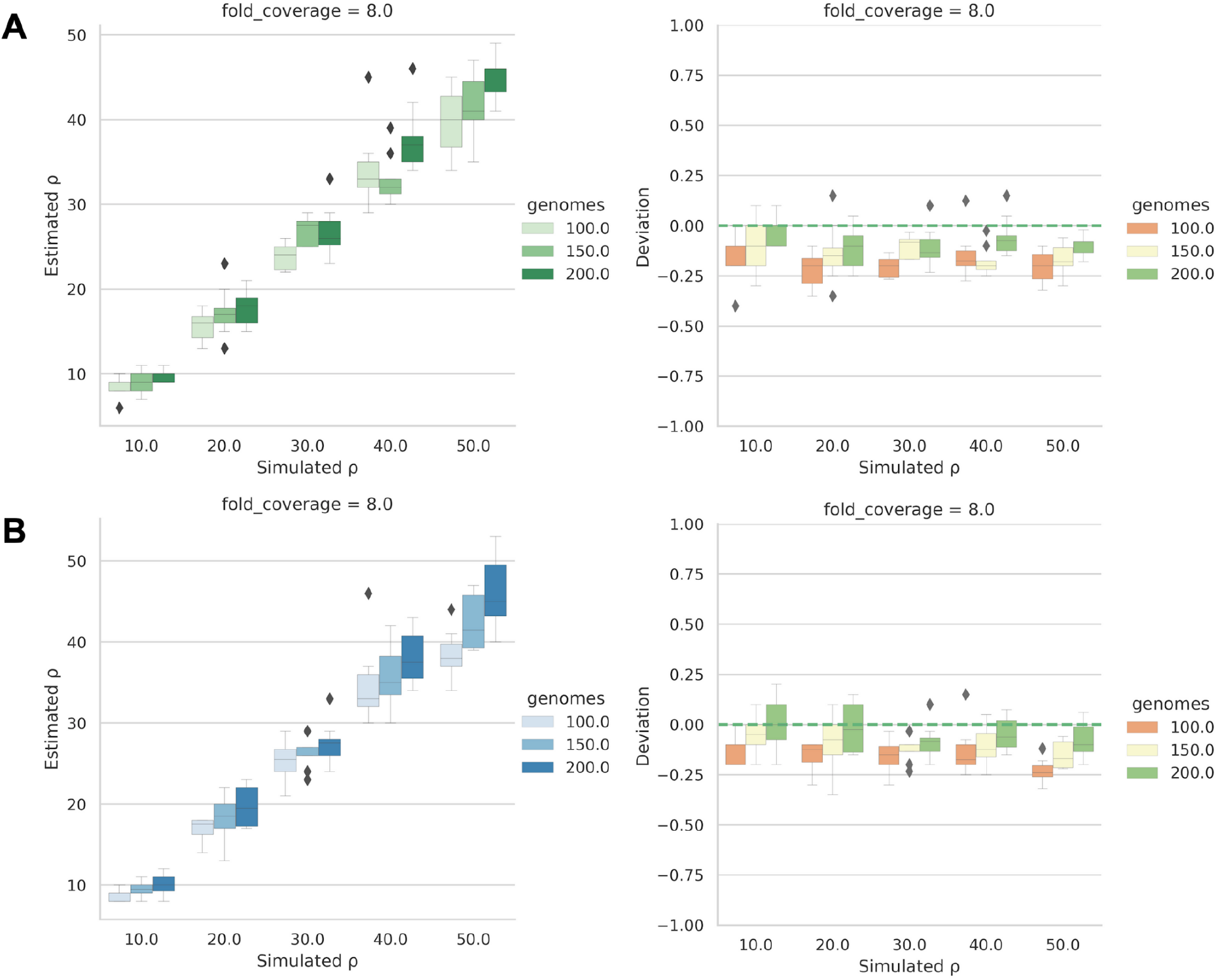
Comparing simulated single end and paired end read datasets in Rhometa. (**A**) Single end results. (**B**) Paired end results

**Figure S3.**
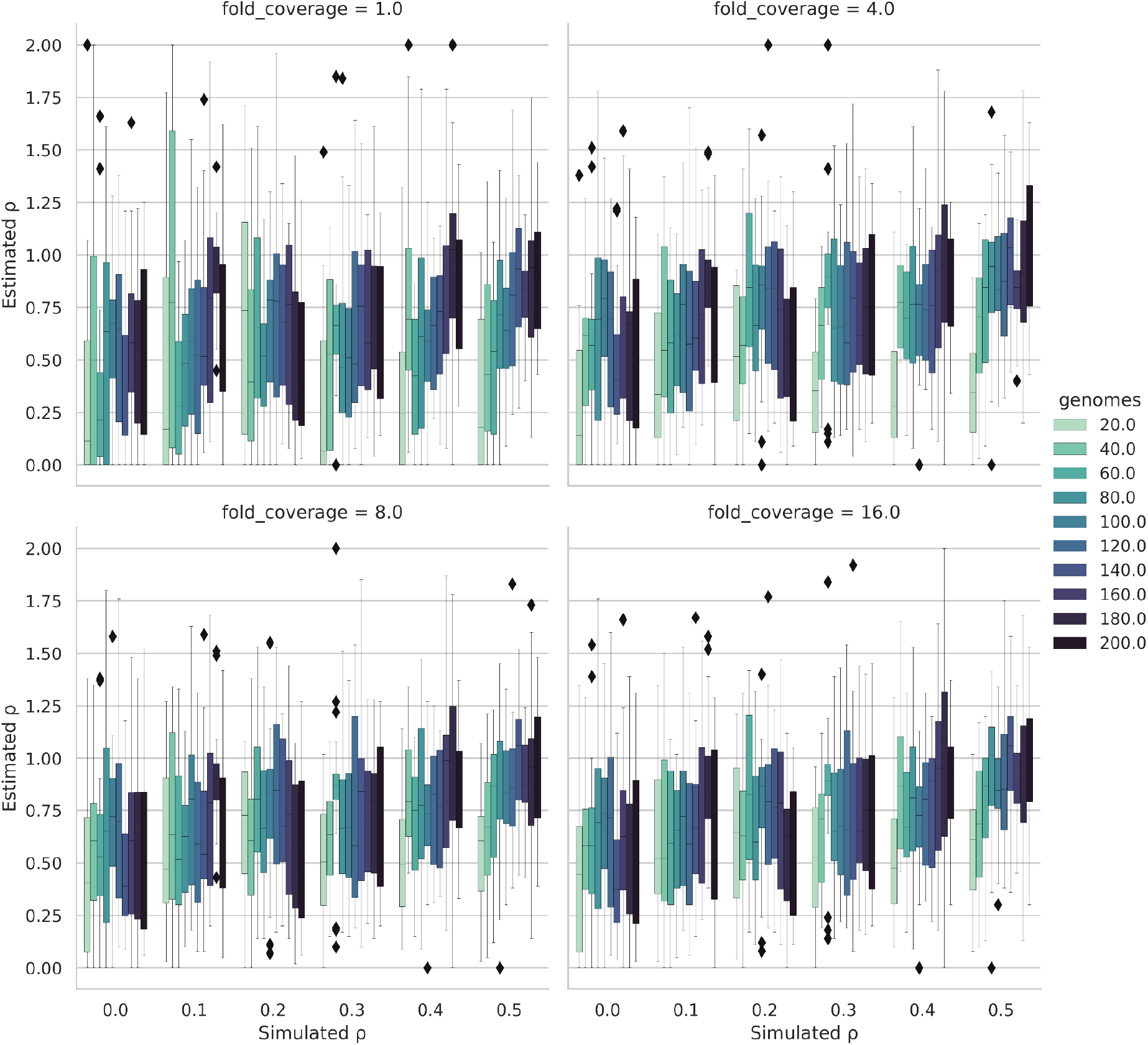
Simulated vs Estimated population recombination rate (ρ) results for Rhometa. Results for varying numbers of simulated genomes and fold coverage values for population recombination rates 0.0, 0.1, 0.2, 0.3, 0.4, 0.5. The simulation parameters used are the same as for population recombination rates [10.0, 20.0, 30.0, 40.0, 50.0], except lookup tables for population recombination rates 0-2 were used (0-2 in 201 steps) for depths of 3-200.

**Figure S4.**
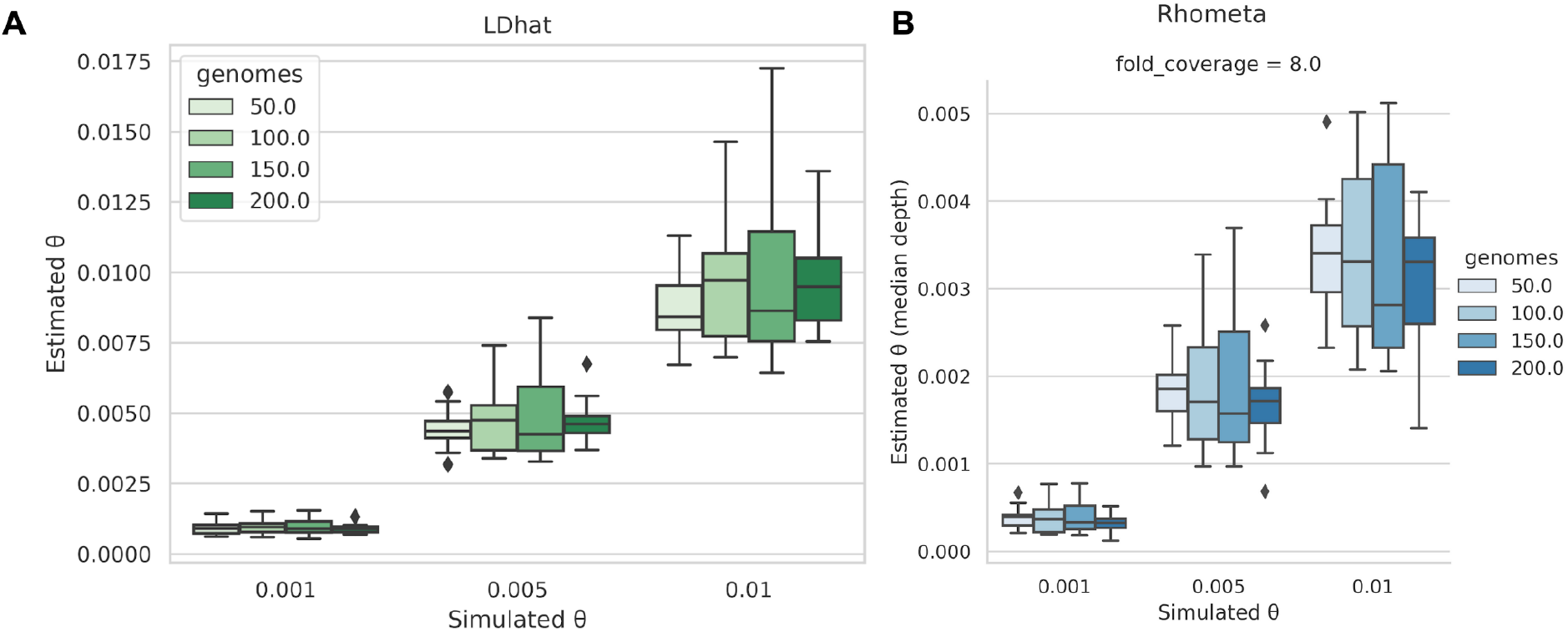
Simulated vs Estimated theta per site (θ) results for LDhat and Rhometa. (**A**) LDhat. Simulated vs Estimated theta per site (**θ**) for varying number of simulated bacterial genomes. (**B**) Rhometa. Simulated vs Estimated theta per site (**θ**) for varying number of simulated bacterial genomes.

## Appendix A

### Lookup configuration

Using the lookup tables requires converting the bi-allelic variant site pairs into a different configuration, which we call the “lookup configuration”. For this step we made use of some functions from Pyrho, where the input has to be in a particular manner, we will explain the LDhat approach first, then the change made for Pyrho.

Let’s consider an earlier example with bi-allelic variant site pair (4,9) with 3 CA and 2 GT as bases, we first need to identify the major allele denoted by 0 and minor allele denoted by 1. For the first site, variant site 4 we have 3 Cs and 2Gs so the major allele (0) is C and the minor allele (1) is G, likewise for the second site variant site 9 we have 3 As and 2 Ts so 0 is A and 1 is T. The lookup configuration is in the format {00, 01, 10, 11}, 00 is site 1 major allele and site 2 major allele, 01 is site 1 major allele and site 2 minor allele and so on.

For 00, the major allele for site 1 is C and the major allele for site 2 is A. Next we look at how many CA pairs there are, we have 3. For 01 we would consider CT and see how many CT pairs we have, which is 0. Continuing in this manner we get the following final results for our bi-allelic pairs, 3 CAs and 2 GTs (CA, CA, CA, GT, GT) becomes {00 : 3, 01 : 0, 10 : 0, 11 : 2}.

### Matching against the lookup table

The index of the lookup tables is in the format {00, 01, 10, 11} by converting the bases for the variant sites into this format we can match against the lookup table and get the likelihoods for each variant site pair, this is done for the entire bi-allelic filtered pairwise table. We used some Pyrho functions for this step, this was done to avoid unnecessary code rewrites and because there are complexities involved in implementation, for instance there needs to be a method in place for handling cases where there is no clear major and minor allele for a site pair i.e. there are a even number of alleles, such as a site pair position with 2 AA and 2 TT. Pyrho, specifically their rho_splines.py script and the compute_splines method therein, has an excellent approach to this step, which we have incorporated into our program. The pyro approach, however, requires the input be formatted in a specific manner.

For the Pyrho method, which is quite a bit simpler, let’s again consider our example with bi-allelic variant site pair (4,9) with 3 CA and 2 GT as bases. First we consider the unique bases at each variant site, for site 1 (variant site 4) we have C and G and for site 2 (variant site 9) we have A and T. For the next step Pyrho encodes the nucleotide bases in this manner { A : 2, C : 3, G : 4, T : 5 }, so for site 1 we have C : 3 and G : 4 and for site 2 we have A : 2 and T : 5. We then use the number associated to the base to determine the major and minor allele per site, for site 1 in G with associated number 4 is the major allele (0) since it is greater than C with associated number 3 which is the minor allele (1), likewise the major and minor alleles for site 2 are T and A respectively.

Having identified the major and minor allele for site 1 and site 2, this needs to be converted in the lookup configuration {00, 01, 10, 11} using the method described earlier the major alleles for site 1 and site 2 are G and T respectively, which is 00, we have 2 GTs so the value for 00 is 2, continuing in this manner we get { 00: 2, 01 : 0, 10 : 0, 11 : 3}. This process is applied to the entire bi-allelic pairwise table, then the lookup configurations from this table along with an appropriate size lookup table can be used by Pyrho to perform the matching and get the likelihoods for the variant site pairs.

